# Fatty acid oxidation participates of the survival to starvation, cell cycle progression and differentiation in the insect stages of *Trypanosoma cruzi*

**DOI:** 10.1101/2021.01.08.425864

**Authors:** Rodolpho Ornitz Oliveira Souza, Flávia Silva Damasceno, Sabrina Marsiccobetre, Marc Biran, Gilson Murata, Rui Curi, Frédéric Bringaud, Ariel Mariano Silber

## Abstract

During its complex life cycle, *Trypanosoma cruzi* colonizes different niches in its insect and mammalian hosts. This characteristic determined the types of parasites that adapted to face challenging environmental cues. The primary environmental challenge, particularly in the insect stages, is poor nutrient availability. These *T. cruzi* stages could be exposed to fatty acids originating from the degradation of the perimicrovillar membrane. In this study, we revisit the metabolic fate of fatty acid breakdown in *T. cruzi*. Herein, we show that during parasite proliferation, the glucose concentration in the medium can regulate the fatty acid metabolism. At the stationary phase, the parasites fully oxidize fatty acids. [U-^14^C]-palmitate can be taken up from the medium, leading to CO_2_ production via beta-oxidation. Lastly, we also show that fatty acids are degraded through beta-oxidation. Additionally, through beta-oxidation, electrons are fed directly to oxidative phosphorylation, and acetyl-CoA is supplied to the tricarboxylic acid cycle, which can be used to feed other anabolic pathways such as the *de novo* biosynthesis of fatty acids.

**Author Summary:** *Trypanosoma cruzi* is a protist parasite with a life cycle involving two types of hosts, a vertebrate one (which includes humans, causing Chagas disease) and an invertebrate one (kissing bugs, which vectorize the infection among mammals). In both hosts, the parasite faces environmental challenges such as sudden changes in the metabolic composition of the medium in which they develop, severe starvation, osmotic stress and redox imbalance, among others. Because kissing bugs feed infrequently in nature, an intriguing aspect of *T. cruzi* biology (it exclusively inhabits the digestive tube of these insects) is how they subsist during long periods of starvation. In this work, we show that this parasite performs a metabolic switch from glucose consumption to lipid oxidation, and it is able to consume lipids and the lipid-derived fatty acids from both internal origins as well as externally supplied compounds. When fatty acid oxidation is chemically inhibited by etomoxir, a very well-known drug that inhibits the translocation of fatty acids into the mitochondria, the proliferative insect stage of the parasites has dramatically diminished survival under severe metabolic stress and its differentiation into its infective forms is impaired. Our findings place fatty acids in the centre of the scene regarding their extraordinary resistance to nutrient-depleted environments.

## Introduction

*T. cruzi*, a flagellated parasite, is the causative agent of Chagas disease, a neglected health problem endemic to the Americas [1]. The parasite life cycle is complex, alternating between replicative and non-replicative forms in two types of hosts, mammalians and triatomine insects [2]. In mammalian hosts, two primary forms are recognized: replicative intracellular amastigotes and nondividing trypomastigotes, which are released from infected host cells into the extracellular medium. After being released from infected cells, trypomastigotes can spread the infection by infecting new cells, or they can be ingested by a triatomine bug during its blood meal. Once inside the invertebrate host, the ingested trypomastigotes differentiate into epimastigotes, which initiate their proliferation and colonization of the insect digestive tract [3]. Once the epimastigotes reach the final portion of the digestive tube, they initiate differentiation into non-proliferative, infective metacyclic trypomastigotes. These forms will be expelled during a new blood meal and will be able to infect a new vertebrate host [2,4–6].

The diversity of environments through which *T. cruzi* passes during its life cycle (i.e., the digestive tube of the insect vector, the bloodstream and the mammalian cells cytoplasm) subjects it to different levels of nutrient availability [3,7]. Therefore, this organism evolved a robust, flexible and efficient metabolism [5,8]. As an example, it was recognized early on that epimastigotes are able to rapidly switch their metabolism, allowing the consumption of carbohydrates and different amino acids [9,10]. Several studies identified aspartate, asparagine, glutamate [11], proline [12–14], histidine [15], alanine [11,16] and glutamine [11,17] as oxidisable energy sources. Despite the quantity of accumulated information on amino acid and carbohydrate consumption, little is known about how *T. cruzi* uses fatty acids and how these compounds contribute to the parasite’s metabolism and survival. In this study, we explore fatty acid metabolism in *T. cruzi.* We also address fatty acid regulation by external glucose levels and the involvement of their oxidation in the replication and differentiation of *T. cruzi* insect stages.

## Methods

### Parasites

Epimastigotes of *T. cruzi* strain CL clone 14 were maintained in the exponential growth phase by sub-culturing them for 48 h in Liver Infusion Tryptose (LIT) medium at 28 °C [18]. Metacyclic trypomastigotes were obtained through the differentiation of epimastigotes at the stationary growth phase in TAU-3AAG (Triatomine Artificial Urine supplemented with 10 mM proline, 50 mM glutamate, 2 mM aspartate and 10 mM glucose) as previously reported [19].

### Fatty acid oxidation assays

#### Preparation of palmitate-BSA conjugates

Sodium palmitate at 70 mM was solubilized in water by heating it up to 70 °C. BSA free fatty acids (FFA BSA) (Sigma®) was dissolved in PBS and warmed up to 37 °C with continuous stirring. Solubilized palmitate was added to BSA at 37 °C with continuous stirring (for a final concentration of 5 mM in 7% BSA). The conjugated palmitate-BSA was aliquoted and stored at −80 °C [20].

#### CO_2_ production from oxidisable carbon sources

To test the production of CO_2_ from palmitate, glucose or histidine, exponentially growing epimastigotes (5×10^7^ mL^−1^) were washed twice in PBS and incubated for different times (0, 30, 60 and 120 min) in the presence of 0.1 mM of palmitate spiked with 0.2 μCi of ^14^C-U-substrates. To trap the produced CO_2_, Whatman paper was embedded in 2 M KOH solution and was placed in the top of the tube. The ^14^CO_2_ trapped by this reaction was quantified by scintillation [15,16].

#### ^1^H-NMR analysis of the exometabolome

Epimastigotes (1×10^8^ mL^−1^) were collected by centrifugation at 1,400 x *g* for 10 min, washed twice with PBS and incubated in 1 mL (single point analysis) of PBS supplemented with 2 g/L NaHCO_3_ (pH 7.4). The cells were maintained for 6 h at 27 °C in incubation buffer containing [U-^13^C]-glucose, non-enriched palmitate or no carbon sources. The integrity of the cells during the incubation was checked by microscopic observation. The supernatant (1 mL) was collected and 50 μl of maleate solution in D_2_O (10 mM) was added as an internal reference. ^1^H-NMR spectra were collected at 500.19 MHz on a Bruker Avance III 500 HD spectrometer equipped with a 5 mm Prodigy cryoprobe. The measurements were recorded at 25 °C. The acquisition conditions were as follows: 90° flip angle, 5,000 Hz spectral width, 32 K memory size, and 9.3 sec total recycling time. The measurements were performed with 64 scans for a total time of close to 10 min and 30 sec. The resonances of the obtained spectra were integrated and the metabolite concentrations were calculated using the ERETIC2 NMR quantification Bruker program.

#### Oxygen consumption

To evaluate the importance of internal fatty acid sources in O_2_ consumption, exponentially growing parasites were treated or not treated with 500 μM ETO (the inhibitor of carnitine palmitoyltransferase 1), washed twice in PBS and resuspended in Mitochondrial Cellular Respiration (MCR) buffer. The rates of oxygen consumption were measured using intact cells in a high-resolution oxygraph (*Oxygraph-2k; Oroboros Instruments*, Innsbruck, Austria). Oligomycin A (0.5 μg/mL) and FCCP (0.5 μM) were sequentially added to measure the optimal non-coupled respiration and the respiration leak state, respectively. The data were recorded and treated using *DatLab 7* software [15,16,21].

### Mitochondrial activity assays

#### MTT and Alamar Blue

The parasites were washed twice and incubated in PBS supplemented with mM palmitate in 0.35% FFA BSA, 0.35% FFA BSA alone, and 5 mM glucose, and 5 mM histidine or not supplemented media were used as controls (positives and negative, respectively). The cell viability was evaluated at 24 h and 48 h after incubation using the MTT assay, as described in [15,16].

#### Alamar Blue

The parasites were washed twice and incubated in PBS or PBS supplemented with 500 μM ETO in 96-well plates. The plates were maintained at 28 °C during all the experiments. After every 24 h, the cells were incubated with 0.125 μg.mL^−1^ of Alamar Blue reagent and kept at 28 °C for 2 h under protection from light. The fluorescence was accessed using the wavelengths λ_exc_ = 530 nm and λ_em_ = 590 nm in the SpectraMax® i3 (Molecular Devices) plate reader.

### Measurement of intracellular ATP content

The intracellular ATP levels were assessed using a luciferase assay kit (Sigma-Aldrich ®), as described in [15–17]. In brief, the parasites were incubated in PBS supplemented (or not) with 0.1 mM palmitate, 0.35% FFA BSA, 5 mM glucose or 5 mM histidine for 24 h at 28 °C. The ATP concentrations were determined by using a calibration curve with ATP disodium salt (Sigma), and the luminescence at 570 nm was measured as indicated by the manufacturer.

### Enzymatic activities

#### Carnitine palmitoyltransferase 1 (CPT1)

The epimastigotes were washed twice in PBS (1,000 × *g*, 5 min at 4 °C), resuspended in buffered Tris-EDTA (100 mM, 2.5 mM and 0.1% Triton X-100) containing 1 μM phenylmethyl-sulphonyl fluoride (PMSF), 0.5 mM N-alpha-p-tosyl-lysyl-chloromethyl ketone (TLCK), 0.01 mg aprotinin and 0.1 mM trans-epoxysuccinyl-L-leucyl amido (4-guanidino) butane (E-64) as a protease inhibitors (Sigma Aldrich®) and lysed by sonication (5 pulses for 1 min each, 20%). The lysates were clarified by centrifugation at 10,000 × *g* for 30 min at 4 °C. The soluble fraction was collected and the proteins were quantified by Bradford method [22] and adjusted to 0.1 mg/mL protein. The reaction mixture contained 0.5 mM L-carnitine, 0.1 mM palmitoyl-CoA and 2.5 mM DTNB in Tris-EDTA buffer (pH = 8.0). The CPT1 activity was measured spectrophotometrically at 412 nm by DTNB reaction with free HS-CoA, forming the TNB^−^ ion. To calculate the specific activity, the absorbance values were converted into molarity by using the TNB^−^ extinction molar coefficient of 12,000 M^−1^.s^−1^ [23]. As a blank, we performed the same assay without adding the substrate. All the enzymatic assays were performed in 96-well plates at a final volume of 0.2 mL in the SpectraMax® i3 (Molecular Devices).

#### Acetyl-CoA carboxylase (ACC)

The ACC activity was measured spectrophotometrically by coupling its enzymatic reaction with that of citrate synthase (CS), which uses oxaloacetate and acetyl-CoA to produce citrate. Measurements were performed at the end-points in two steps. First, the reaction mixture contained 100 mM potassium phosphate buffer (pH = 8.0), 15 mM KHCO_3_, 5 mM MnCl_2_, 5 mM ATP, 1 mM acetyl-CoA and 0.1 μM biotin. The reaction was initiated by adding 0.1 mg of cell extract and developed using 15 min incubations at 28 °C. The reaction was stopped by adding perchloric acid 40% (v/v) and centrifuged 10,000 × *g* for 15 min at 4 °C. The second reaction was performed by using 0.1 mL of the supernatant from the first reaction, 20 mM oxaloacetate and 0.5 mM of DTNB in 100 mM potassium phosphate buffer (pH = 8.0). The reaction was initiated by adding 0.5 units of CS (Sigma Aldrich©). To calculate the specific activity of ACC, we converted the absorbance values to molarity by using the TNB^−^ extinction molar coefficient of 12,000 M^−1^.s^−1^. For the blank reaction, we performed the same assay without acetyl-CoA [24].

#### Hexokinase (HK)

The HK activity was measured as described in [25]. Briefly, the activity was measured by coupling the hexokinase activity with a commercial glucose-6-phosphate dehydrogenase, which oxidizes the glucose-6-phosphate (G6PD, SIGMA) resulting from the HK activity with the concomitant reduction of NADP^+^ to NADPH. The resulting NADPH was spectrophotometrically monitored at 340 nm. The reaction mixture contained 50 mM Triethanolamine buffer pH 7.5, 5 mM MgCl_2_, 100 mM KCl, 10 mM glucose, 5 mM ATP and 5 U of commercial G6PD. To calculate the specific activity, the absorbance values were converted to molarity using the NADP(H) extinction molar coefficient of 6,220 M^−1^.s^−1^.

#### Serine palmitoyltransferase (SPT)

The SPT activity was measured through the reduction of the DTNB reaction by the free HS-CoA, forming the TNB^−^ ion, which was measured spectrophotometrically at 412 nm as previously described [23]. In brief, the epimastigotes were washed twice in PBS, resuspended in Tris-EDTA buffer (100 mM/2.5 mM) containing Triton X-100 0.1% and lysed by sonication (20% of potency, during 2 min). The reaction mixture contained 0.1 mg of protein free-cell extract, 0.5 mM L-serine, 0.1 mM palmitoyl-CoA and 2.5 mM DTNB in Tris-EDTA buffer (100 mM/2.5 mM) pH = 8.0 [26]. To calculate the specific activity, we converted the absorbance values to molarity using the TNB^−^ extinction molar coefficient of 12,000 M^−1^.s^−1^. For the blank reaction, we performed the same assay without adding palmitoyl-CoA. All the enzymatic assays were performed in 96-well plates in a final volume of 0.2 mL in the SpectraMax® i3 (Molecular Devices).

### Glucose and triglyceride quantification

Spent LIT medium from epimastigote cultures was collected by recovering the supernatants from a centrifugation (10,000 × *g* for 15 min at 4 °C). Each sample of spent LIT was analysed for its glucose and triglyceride contents using commercial kits (triglyceride monoreagent and glucose monoreagent by Bioclin Brazil) according to the manufacturer’s instructions. These kits are based on colorimetric enzymatic reactions, and the absorbance of each assay was measured in 96-well plates at a final volume of 0.2 mL in the SpectraMax® i3 (Molecular Devices).

### Proliferation assays

Exponentially growing *T. cruzi* epimastigotes (5×10^7^ mL^−1^) were treated with different concentrations of ETO or not treated (negative control) in LIT medium. As a positive control for growth inhibition, we used a combination of rotenone (60 μM) and antimycin (0.5 μM) [27]. The parasites (2.5×10^6^ mL^−1^) were transferred to 96-well plates and then incubated at 28 °C. The cell proliferation was quantified by reading the optical density (OD) at 620 nm for eight days. The OD values were converted to cell numbers using a linear regression equation previously obtained under the same conditions. Each experiment was performed in quadruplicate [28].

### Flow cytometry analyses

#### Cell death

Epimastigotes in the exponential phase of growth were maintained in LIT and treated with ETO 500 μM for 5 days. After the incubation time, the parasites were analysed as described in [28]. The cells were analysed by flow cytometry (FACScalibur BD Biosciences).

#### Cell cycle (DNA content)

Epimastigotes in the exponential phase of growth were maintained in LIT and treated with ETO 500 μM over 5 days. After the incubation time, the parasites were washed twice in PBS and resuspended in lysis buffer (phosphate buffer Na_2_HPO_4_ 7.7 mM; KH_2_PO_4_ 2.3 mM; pH = 7.4) and digitonin 64 μM. After incubating on ice for 30 min, propidium iodide 0.2 μg/mL was added. The samples were analysed by flow cytometry (Guava) adapted from [29].

#### Fatty acid staining using BODIPY® 500/510

Exponentially growing epimastigotes were kept in LIT medium to reach three different cell densities (2.5×10^7^ mL^−1^, 5×10^7^ mL^−1^ and 10^8^ mL^−1^) in 24-well plates at 28 °C. Twenty-four hours before the flow cytometry analysis, the parasites were treated with 1 μM C_1_-BODIPY® 500/510-C_12_. This fluorophore allows for measurements of the relationship between fatty acid accumulation and consumption by shifting the fluorescence filter. The samples were collected, washed twice in PBS and incubated in 4% paraformaldehyde for 15 min. After incubation, the cells were washed twice with PBS and suspended in the same buffer. Flow cytometry analysis was performed with FL-1 and FL-2 filters in a FACS Fortessa DB®. The results were analysed using FloJo software.

### Fluorescence microscopy

The parasites were maintained in LIT medium as previously reported for fatty acid staining using *BODIPY® 500/510*. After incubation, the cells were washed twice in PBS and placed on glass slides. The images were acquired with a digital DFC 365 FX camera coupled to a DMI6000B/AF6000 microscope (Leica). The images were analysed using ImageJ software.

## Results

### Palmitate supports ATP synthesis in *T. cruzi*

We initially investigated the ability of *T. cruzi* epimastigotes to oxidize fatty acids. To this end, we used palmitate as a proxy for fatty acids in general. The parasites were incubated with 0.1 mM ^14^C-[U]-palmitate, which allowed us to measure the production of 1.3 nmoles of CO_2_ derived from palmitate oxidation during the first 60 min and 1.5 extra nmoles during the following 60 min (Fig 1a). This finding indicated that beta-oxidation and the further ‘burning’ of the resulting acetyl-CoA is operative in epimastigote mitochondria. Because palmitate is taken up from extracellular medium and oxidized to CO_2_, it is reasonable to assume that it could contribute to resistance to severe nutritional stress. To support this idea, we tested the ability of palmitate to extend parasite survival under extreme nutritional stress. Parasites were incubated for 24 and 48 h in PBS (negative control, in this condition we expected the lower viability after the incubations), 0.1 mM palmitate in PBS supplemented with BSA (as a palmitate carrier), 5.0 mM histidine in PBS or 5.0 mM glucose in PBS (both positive controls, since it is well knowing the ability of both metabolites to extend the parasites’ viability in metabolic stress conditions, see [15]). As an additional negative control, we used PBS supplemented with BSA without added palmitate. The viability of these cells was assayed by measuring the total reductive activity by MTT assay. Additionally, we measured the total ATP levels. Cells incubated in the presence of palmitate showed higher viability than the negative controls, but not as high as that of parasites incubated with glucose or histidine (Fig 1b). Consistently, parasites incubated in the presence of palmitate showed higher ATP contents than both negative controls. However, the intracellular ATP levels in the cells incubated with palmitate were diminished by half when compared to parasites incubated with histidine. Interestingly, the palmitate kept the ATP content at levels comparable to glucose (Fig 1c).

**Figure 1.**
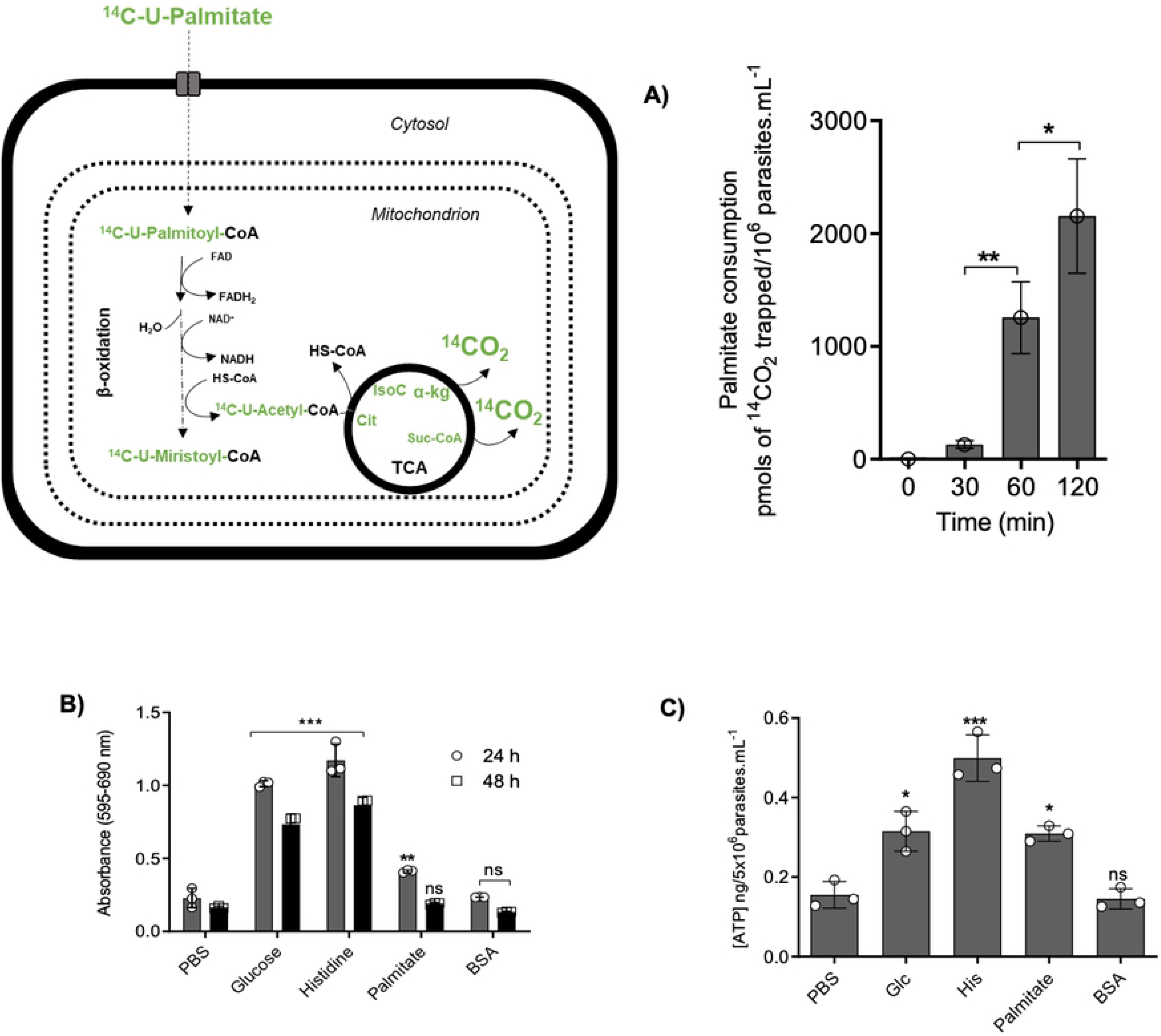
Palmitate oxidation promotes ATP production and viability in epimastigote forms under starvation. Schematic representation of ^14^C-U-palmitate metabolism. The metabolites corresponding to labelled palmitate metabolism are presented in green. A) ^14^CO_2_ production from epimastigotes incubated in PBS with ^14^C-U-palmitate 100 μM. The ^14^CO_2_ was captured at 0, 30, 60 and 120 min. B) Viability of epimastigote forms after incubation with different carbon sources and palmitate. The viability was assessed after 24 and 48 h by MTT assay. C) The intracellular ATP content was evaluated following incubation with different energy substrates or not (PBS, negative control). The ATP concentration was determined by luciferase assay and the data were adjusted by the number of cells. A statistical analysis was performed with one-way ANOVA followed by Tukey’s post-test at p < 0.05 using the GraphPad Prism 8.0.2 software program. We represent the level of statistical significance in this figure as follows: *** p value < 0.001; ** p value < 0.01; * p value < 0.05. For a p value > 0.05 we consider the differences to be not significant (ns).

### Epimastigote forms excrete acetate as a primary end-product of palmitate oxidation

Because the epimastigotes were able to oxidize ^14^C-U-palmitate to ^14^CO_2_, we were interested in analysing their exometabolome and comparing it with that of parasites exclusively consuming glucose, palmitate or without any carbon source. Thus, we subjected exponentially growing parasites to 16 h of starvation and then incubated them for 6 h in the presence of 0.3 mM palmitate, 10 mM ^13^C-U-glucose or without any carbon source. For the control, we analysed a sample of non-starved parasites. After the incubations, the extracellular media were collected and analysed by ^1^H-NMR spectrometry. As expected, all the incubation conditions produced different flux profiles for excreted metabolites (Fig 2 and S1 Fig). Under our experimental conditions, the non-starved parasites primarily excreted succinate and acetate in similar quantities, and alanine and lactate to a lesser extent. Parasites starved for 16 h in PBS and left to incubate in the absence of other metabolites had diminished succinate production (~7-fold) but increased acetate production three-fold compared to the non-starved parasites. It is relevant to stress that the only possible origin for these metabolites are internal carbon sources (ICS). Notably, no other excreted metabolites were detected under these conditions, indicating that under starvation, most of the ICS are transformed into acetate as an end product, which is compatible with the oxidation of internal fatty acids. These results raise the question about the metabolic fates of glucose or fatty acids in previously starved parasites. Starved epimastigotes that recovered in the presence of glucose exhibited a profuse excretion of succinate (450-fold the quantity excreted by the starved cells) and roughly equivalent quantities of acetate compared with the starved cells. Interestingly, lactate and alanine were also excreted at similar levels. As expected, the recovery with glucose produced an increase in all the secreted metabolites. However, analysing their distribution is a reconfiguration of the metabolism towards a majority production of succinate. Finally, in epimastigotes incubated with palmitate, we observed an increase in the acetate and alanine production of approximately 2.5 times to the levels in parasites that recovered in the presence of glucose. Interestingly, succinate is excreted in a smaller quantity than acetate and alanine, but still at 10-fold the rate observed in the starved non-recovered cells. Surprisingly, there was also a significant production of pyruvate (not previously described in the literature, and not observed under any other conditions) and a small amount of lactate derived from palmitate.

**Figure 2.**
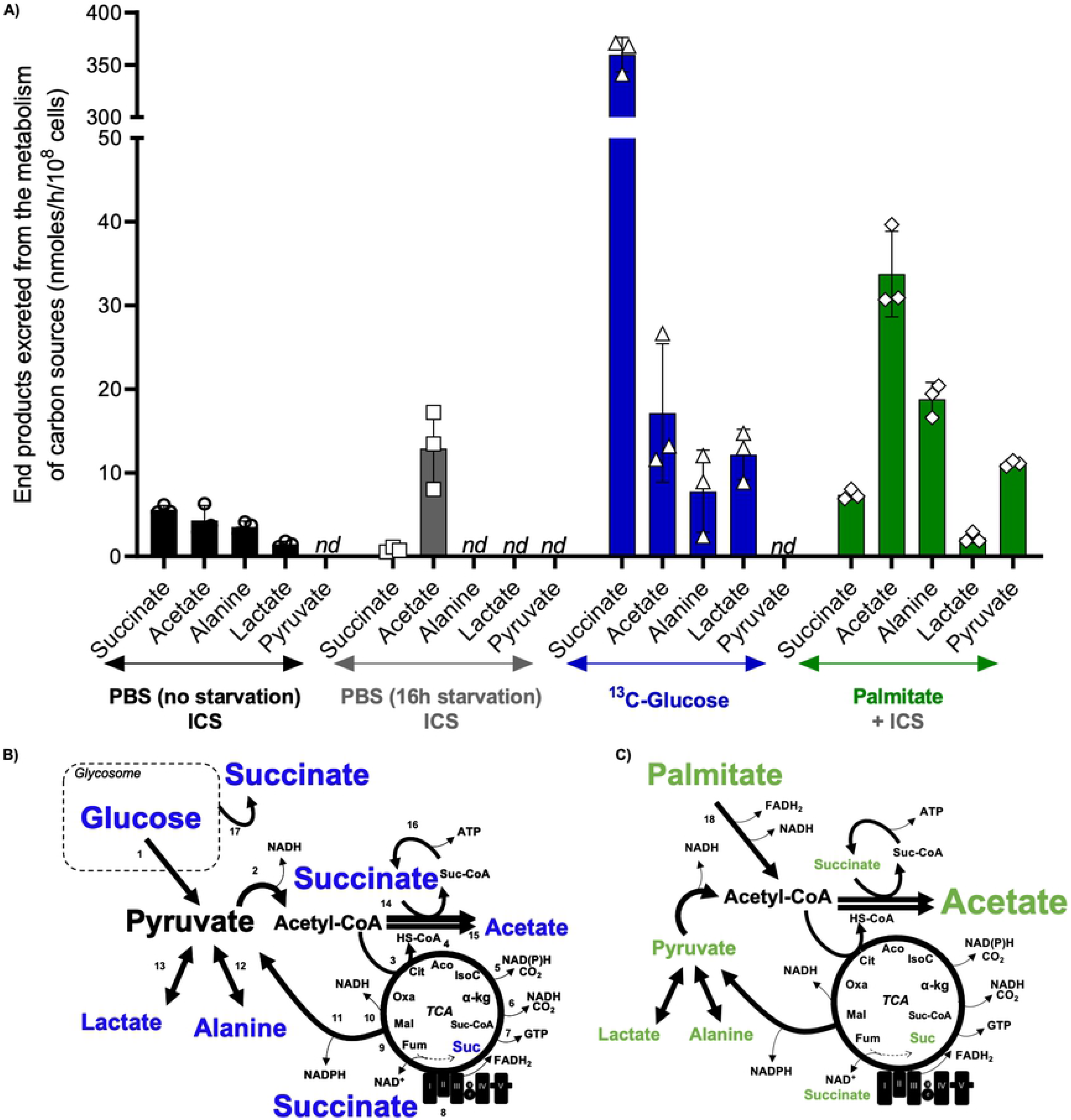
Excreted end products of glucose and palmitate metabolism in epimastigote forms of *T. cruzi*. A) The extracellular medium of epimastigote forms incubated under different conditions was analysed by ^1^H-NMR spectrometry to detect and quantify the end-products. The resulting data were expressed in nmoles/h/10^8^ cells. Means ± SD of three independent experiments. ICS is internal carbon sources; *nd* is non-detectable. B) and C) Schematic representation of the contribution of glucose and palmitate to the metabolism of epimastigote forms of *T. cruzi*. The glycosomal compartment and TCA cycle are indicated. The amount of end-product determined by the font size. Numbers indicates enzymatic steps. 1. Glycolysis; 2. pyruvate dehydrogenase; 3. citrate synthase; 4. aconitase; 5. isocitrate dehydrogenase; 6. α-ketoglutarate dehydrogenase; 7. succinyl-CoA synthetase; 8. Succinate dehydrogenase/complex II/fumarate reductase NADH-dependent; 9. fumarate hydratase; 10. malate dehydrogenase; 11. Malic enzyme; 12. alanine dehydrogenase/alanine aminotransferase; 13. lactate dehydrogenase; 14. acetate:succinyl-CoA transferase; 15. acetyl-CoA hydrolase; 16. succinyl-CoA synthetase; 17. Glycosomal fumarate reductase and 18. Palmitate oxidation by beta-oxidation, resulting in FADH_2_, NADH and acetyl-CoA; Abbreviations: Cit: Citrate, Aco: Aconitate, IsoC: Isocitrate, α-kg: α-Ketoglutarate, Suc-CoA: Succinyl-CoA, Suc: Succinate, Fum: Fumarate, Mal: Malate, and Oxa: Oxaloacetate.

### Glucose metabolism represses the fatty acid oxidation in epimastigotes

Glucose is the primary carbon source for exponentially proliferating epimastigotes, and after its exhaustion from the culture medium, the parasites change their metabolism to use amino acids as carbon sources preferentially [10]. Therefore, we were interested in analysing if this preference for glucose is maintained in relation to the consumption of lipids. To determine if glucose metabolism interferes with the consumption of fatty acids, we created a 48 h proliferation curve using parasites with an initial concentration adjusted to 2.5 × 10^7^ mL^−1^ and quantified them for 24 h each. Under these conditions, the parasites from the beginning of the experiment, at 0 h, are at mid-exponential phase, they are at late exponential phase at 24 h, and at 48 h they reached stationary phase at a concentration of 10 × 10^7^ mL^−1^ (Fig 3A). At 0 h, 24 h and 48 h, the culture medium was collected to measure the remaining glucose and triacylglycerol (TAGs) concentrations (Figs 3B and 3C). Most of the glucose was consumed during the first 24 h (during proliferation), while the concentration of TAGs remained the same. After 48 h of proliferation (stationary phase), the TAG levels and lipid contents of the droplets were decreased by 1.5-fold and 2-fold, respectively, suggesting that glucose is preferentially consumed relative to fatty acids. These data show a decrease in the extracellular TAGs between 24 and 48 h, while the glucose was already almost entirely consumed, suggesting that glucose is negatively regulating the fatty acid catabolism.

**Figure 3.**
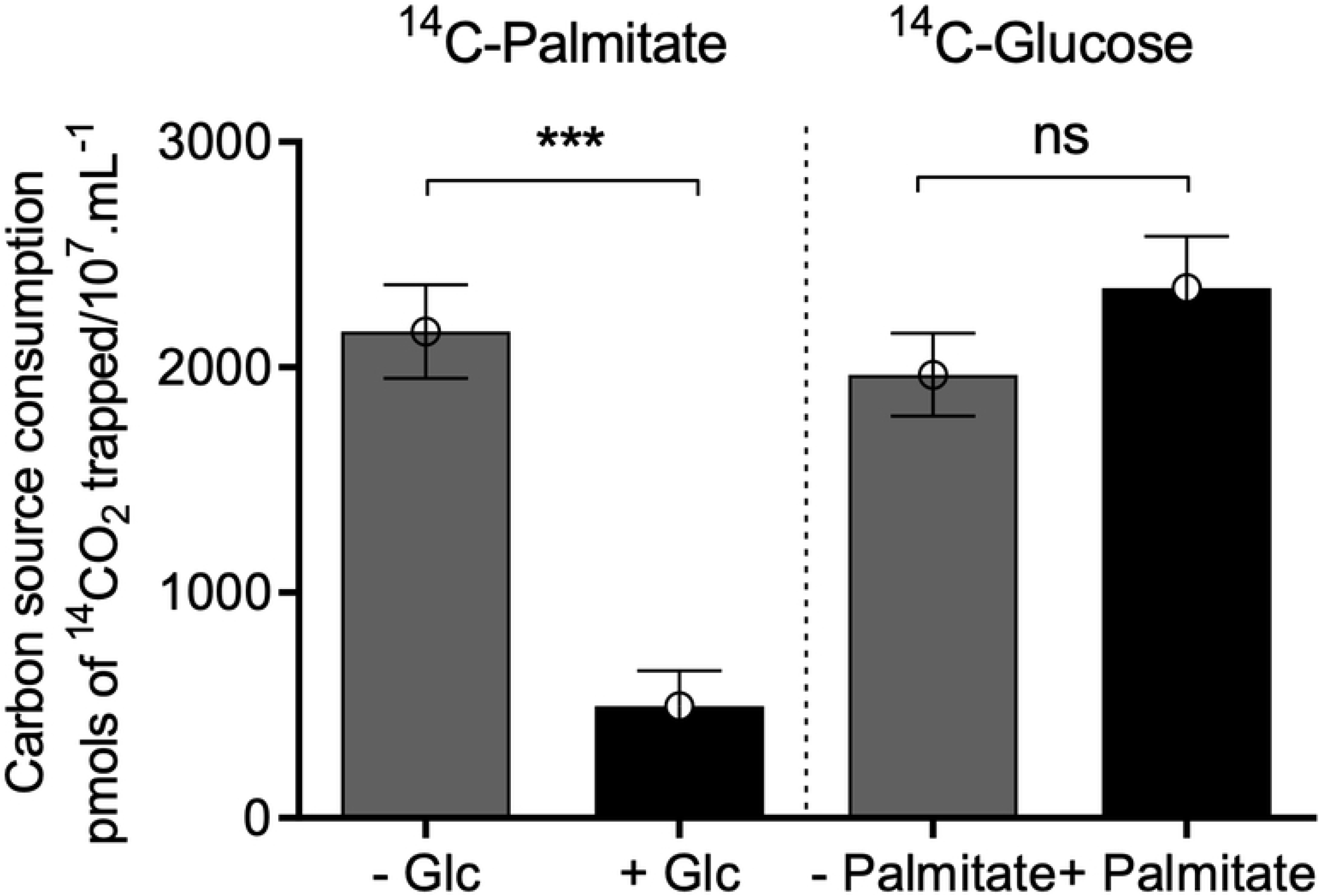
Changes in glucose and triacylglycerol contents in LIT medium. A) Growth curve of epimastigote forms. B) Glucose quantification over 48 h. C) Triacylglycerol levels over 48 h. In each experiment, we collected each medium at different times and subjected it to quantification according to the manufacturer’s instructions. All the experiments were performed in triplicates. Statistical analysis was performed with one-way ANOVA followed by Tukey’s post-test p < 0.05 using the GraphPad Prism 8.0.2 software program. We represent the levels of statistical significance in this figure as follows: *** p value < 0.001; ** p value < 0.01; and * p value < 0.05. For p value > 0.05, we consider the differences not significant (ns).

### Epimastigote forms use endogenous fatty acids to support growth after glucose exhaustion

From the previous results, we learned that under glucose deprivation, TAGs are taken up by the epimastigotes, and internally stored fatty acids are mobilized. However, to date, we did not provide any evidence pointing to their use as reduced carbon sources. To confirm this idea, exponentially proliferating epimastigotes were incubated in PBS supplemented with palmitate and ^14^C-U-glucose, or reciprocally, glucose and ^14^C-U-palmitate. In both cases, the production of ^14^C-labelled CO_2_ was quantified. The presence of 5 mM glucose diminished the release of ^14^CO_2_ from ^14^C-U-palmitate by 90% while the presence of palmitate did not interfere with the production of ^14^CO_2_ from ^14^C-U-glucose (Fig 4). Taken together, our results show that glucose inhibits TAGs and fatty acid consumption, and after glucose exhaustion, a metabolic switch occurs towards the oxidation of internally stored fatty acids.

**Figure 4.**
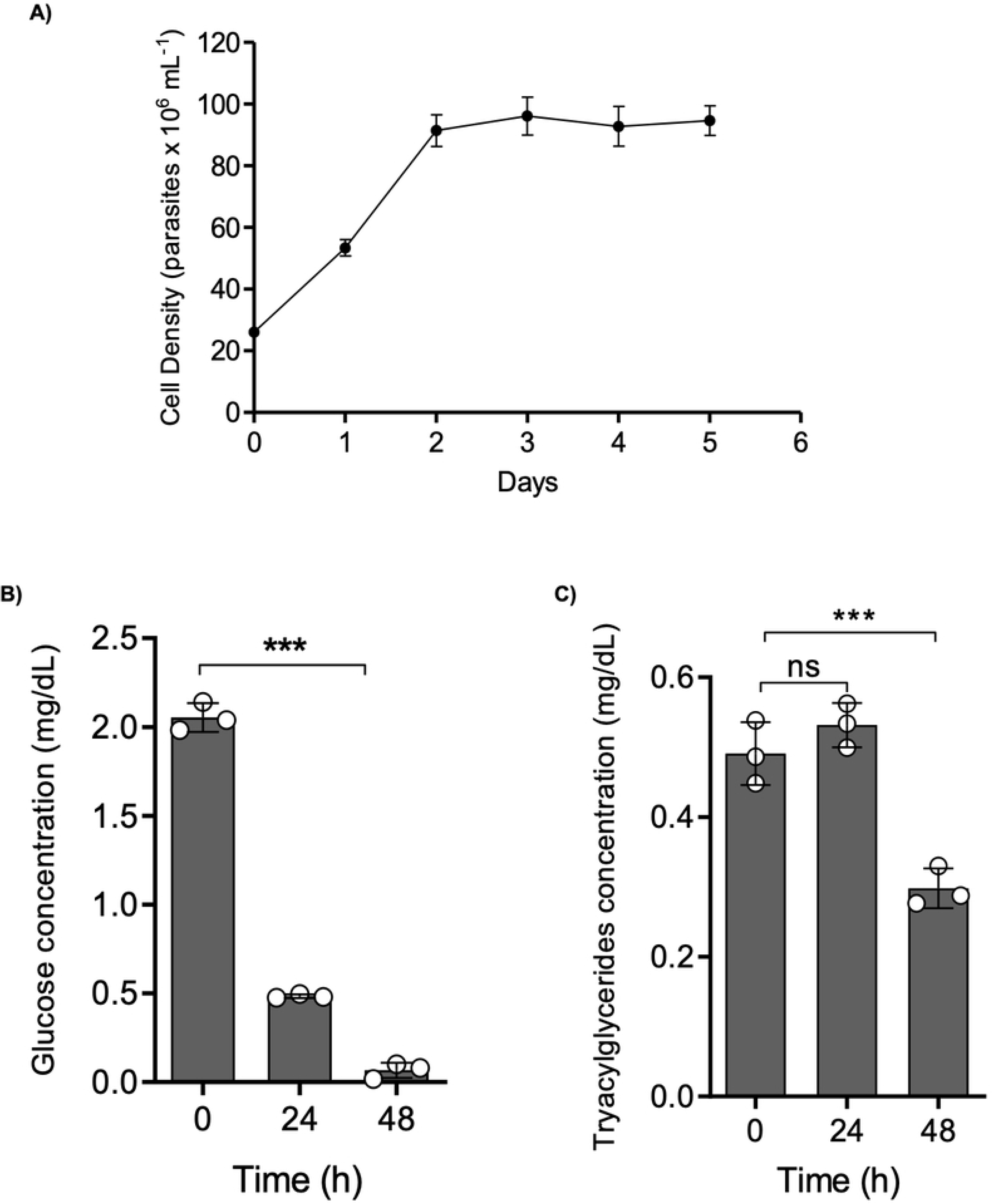
Glucose metabolism inhibits FAO. Parasites were incubated in the presence of ^14^C-U-palmitate + 5 mM glucose and ^14^C-U-glucose + 0.1 mM palmitate in PBS. ^14^CO_2_ production from epimastigotes incubated in PBS. The ^14^CO_2_ was captured after 120 min of incubation. The experiments were performed in triplicates. Statistical analysis was performed with one-way ANOVA followed by Tukey’s post-test p < 0.05 using the GraphPad Prism 8.0.2 software program. We represent the level of statistical significance in this figure as follows: *** p value < 0.001; ** p value < 0.01; and * p value < 0.05. For p value > 0.05, we consider the differences not significant (ns).

To monitor the dynamics of use or accumulation of fatty acids in lipid droplets, we used as a probe a fluorescent fatty acid analogue called BODIPY 500/510 C_1_-C_12_. BODIPY shifts its fluorescence from red to green upon the uptake and catabolism of fatty acids, and from green to red when fatty acids are accumulated in the lipid droplets. Parasites collected at the mid and late exponential proliferation phases and the stationary phase were incubated with 1 μM BODIPY 500/510 C_1_-C_12_ for 16 h, before fluorescence determination by flow cytometry (Figs 5A, 5B and 5C). The fluorescence values increased with the harvesting time (and therefore, with the glucose depletion), indicating the increased uptake and use of fatty acids as substrates by a fatty acyl-CoA synthetase. These data were confirmed by fluorescence microscopy (Fig 5D). Interestingly, parasites in stationary phase showed an accumulation of activated fatty acids in spots along the cell. However, the number of lipid droplets increased upon parasite proliferation (Figs 6A, 6B 6C). This observation indicates that not only fatty acids metabolism is activated after glucose exhaustion, but also the parasite storage of fatty acids into lipid droplets.

**Figure 5.**
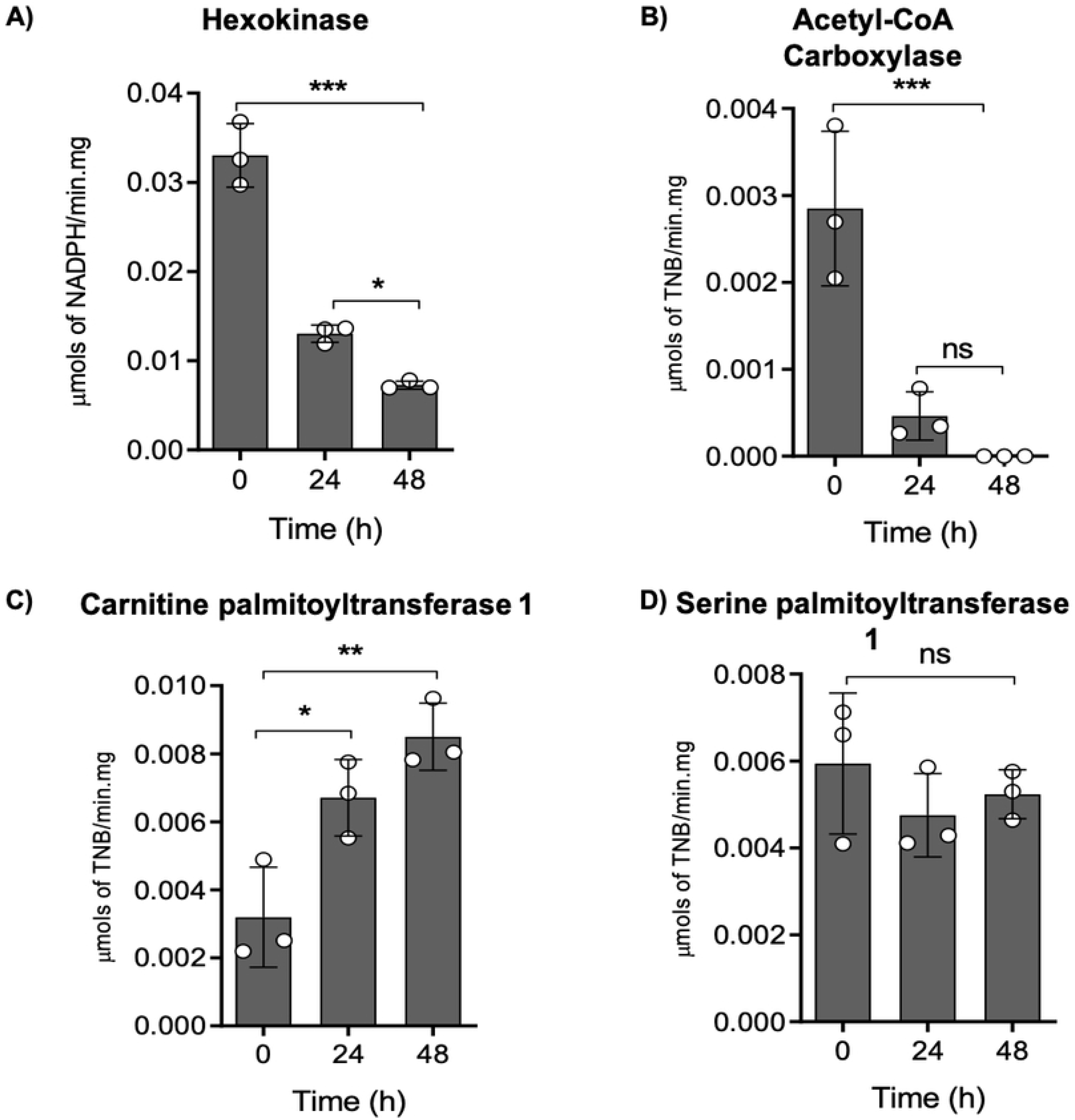
Flow cytometry reveals distinct patterns in fatty acid pools during epimastigote growth. The epimastigotes were treated with 1 μM of BODIPY C_1_-C_12_ (500/510) and analysed by flow cytometry and fluorescence microscopy. A) 0 h. B) 24 h. C) 48 h. In the flow cytometry histograms, dashed peaks represent unstained parasites. Green-filled peaks represent stained parasites. D) Mean fluorescence per cell. The fluorescence for each cell was calculated using ImageJ software. All the experiments were performed in triplicates. Statistical analysis was performed with one-way ANOVA followed by Tukey’s post-test p < 0.05 using the GraphPad Prism 8.0.2 software program. We represent the level of statistical significance in this figure as follows: *** p value < 0.001; ** p value < 0.01; and * p value < 0.05. For p value > 0.05, we consider the differences not significant (ns).

**Figure 6.**
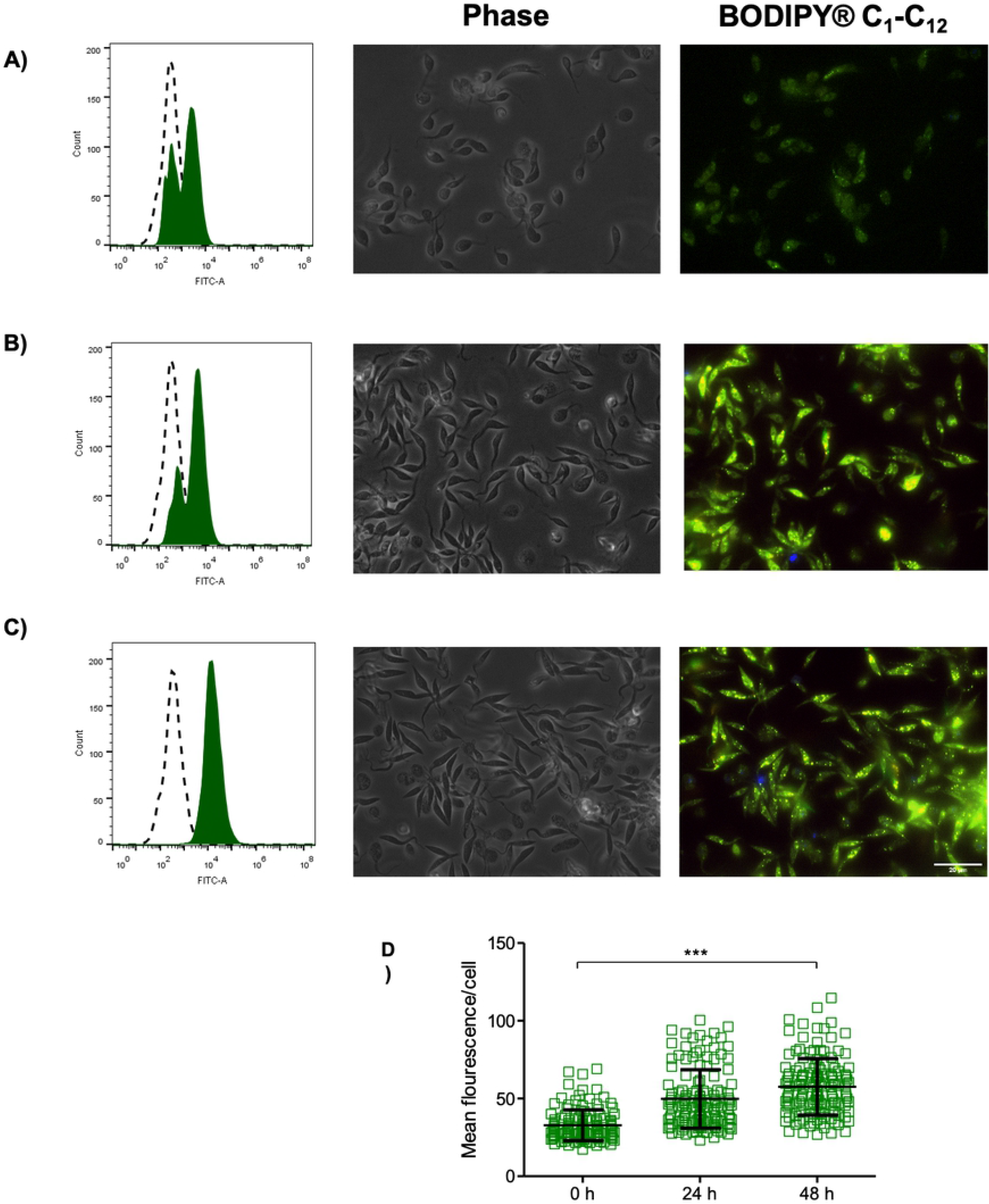
Activities of enzymes related to lipid and glucose metabolism during *T. cruzi* growth curves. A) (HK) Hexokinase B) (ACC) acetyl-CoA carboxylase, C) (CPT1) carnitine-palmitoyltransferase, and D) (SPT) serine palmitoyltransferase. All these activities were measured in crude extracts from epimastigote forms at different moments of the growth curve. All the experiments were performed in triplicates. Time course activities and controls shown in Fig S2. Statistical analysis was performed with one-way ANOVA followed by Tukey’s post-test at p < 0.05, using the GraphPad Prism 8.0.2 software program. We represent the level of statistical significance in this figure as follows: *** p value < 0.001; ** p value < 0.01; and * p value < 0.05. For p value > 0.05 we consider the differences not significant (ns).

To find if the increase in fatty acid pools is accompanied by a change in the levels of enzymes related to fatty acid metabolism, we evaluated the specific activities of the enzymes hexokinase (HK), which is responsible for the initial step of glycolysis and an indicator of active glycolysis; acetyl-CoA carboxylase (ACC), which produces malonyl-CoA for fatty acid synthesis and carnitine palmitoyltransferase 1 (CPT1), the complex that plays a central role in fatty acid oxidation (FAO) by controlling the entrance of long-chain fatty acids into the mitochondria [30]. For the control, we selected the enzyme serine palmitoyltransferase 1 (SPT1), a constitutively expressed protein in *T. cruzi* [31] (Fig 7). The hexokinase activity diminished up to 30% with the progression of the proliferation curve and the correlated depletion of glucose (Fig 7A). In addition, the ACC activity is no more detectable in the stationary phase cells (Fig 7B). By contrast, the CPT1 activity is increased by ~4-fold when the stationary phase is reached (Fig 7C), which confirms that fatty acid degradation occurs in the absence of glucose. It is noteworthy that the high levels of ACC activity in the presence of glucose supports the idea that under these conditions, fatty acids are probably synthesized instead of being catabolized. As expected, SPT1 did not change during the analysed time frame (Fig 7D).

**Figure 6.**
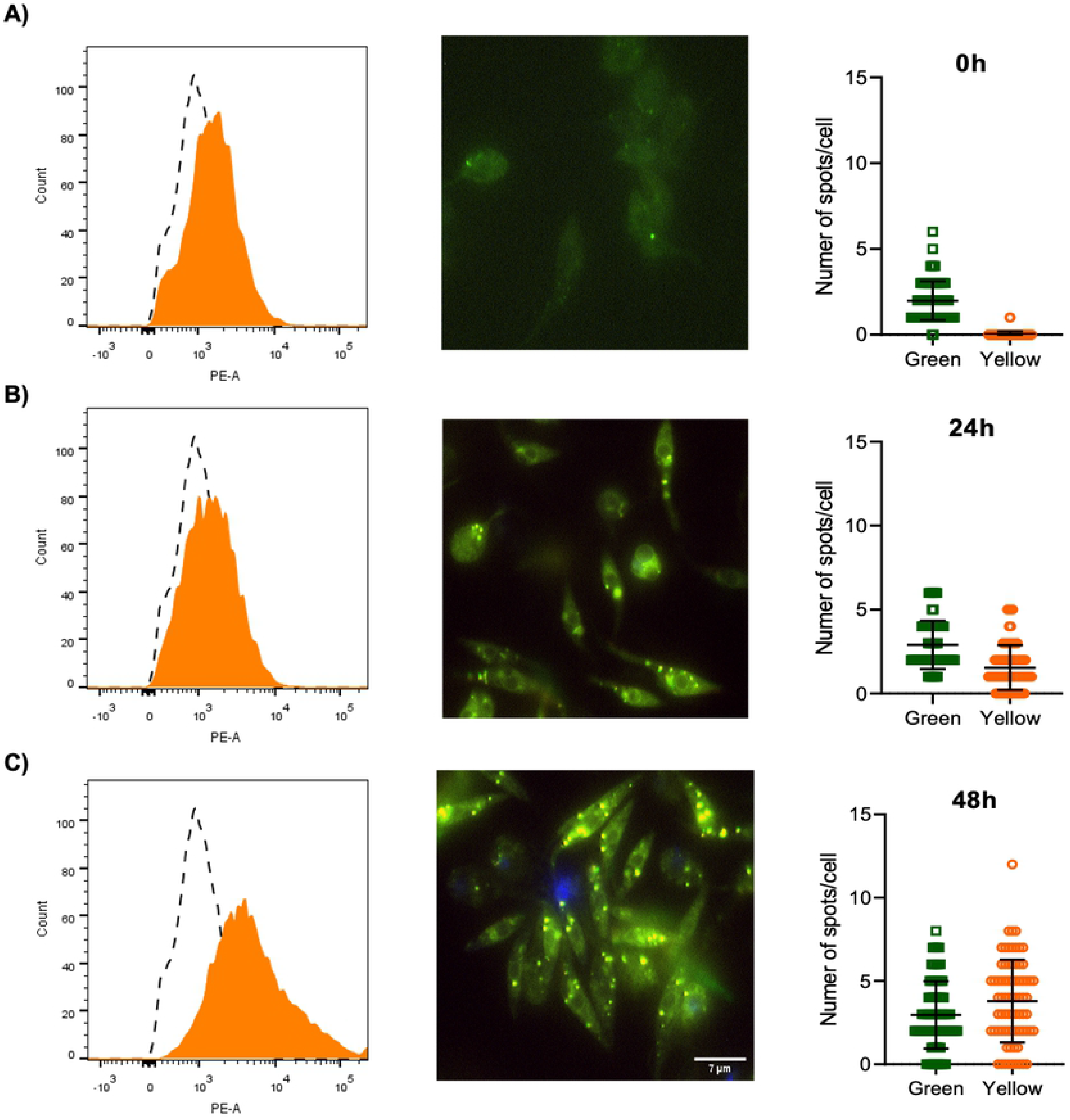
Epimastigote forms accumulates fatty acids into lipid droplets during growth. The epimastigotes were treated with 1 μM BODIPY C_1_-C_12_ (500/510) and analysed by flow cytometry and fluorescence microscopy. A) 0 h. B) 24 h. C) 48 h. In the flow cytometry histograms, dashed peaks represent unstained parasites. Yellow filled peaks represent positively stained parasites. The number of green/yellow spots for each cell was calculated using ImageJ software. All the experiments were performed in triplicates.

### Etomoxir, a CPT1 inhibitor, affects *T. cruzi* proliferation and mitochondrial activity

To investigate the role of FAO in *T. cruzi*, we tested the effect of a well characterized inhibitor of CPT1, etomoxir (ETO), on the proliferation of epimastigotes. Among the ETO concentrations tested here (from 0.1 to 500 μM), only the higher concentration arrested parasite proliferation (Fig 8A). Importantly, the ETO effect was manifested when the parasites reached the late exponential phase (a cell density of approximately 5×10^7^ mL^−1^). This result is consistent with our previous findings showing that FAO (and thus CPT1 activity) acquires an important role at this point in the proliferation curve. To confirm that CPT1 is in fact a target of ETO in *T. cruzi*, we assayed the drug’s effect on the enzyme activity in free cell extracts. Our results showed that 500 μM ETO diminished the CPT1 activity by almost 80% (Fig 8B). To confirm the interference of ETO with the beta-oxidation of fatty acids, parasites incubated in PBS containing ^14^C-U-palmitate were treated with 500 μM ETO to compare their production of ^14^CO_2_ with that of the untreated controls. Palmitate-derived CO_2_ production diminished by 80% in ETO-treated cells compared to untreated parasites (Fig 8C). In addition, ETO treatment did not affect the metabolism of ^14^C-U-glucose or ^14^C-U-histidine, ruling out a possible unspecific reaction of this drug with CoA-SH as described by [32]. Other compounds described as FAO inhibitors were also tested, but none of them inhibited epimastigote proliferation or ^14^CO_2_ production from ^14^C-U-palmitate (S3 Fig). In addition, the BODIPY cytometric analysis of cells treated with 500 μM ETO showed a strong decrease in the CoA acylation levels (activation of fatty acids) with respect to the untreated controls (Fig 8D), as confirmed by fluorescence microscopy (Fig 8D). To reinforce the validation of ETO for further experiments, a set of controls are offered in S3 Fig. Our preliminary conclusion is that ETO inhibited beta-oxidation by inhibiting CPT1, confirming that the breakdown of fatty acids is important to proliferation progression in the absence of glucose.

**Figure 8.**
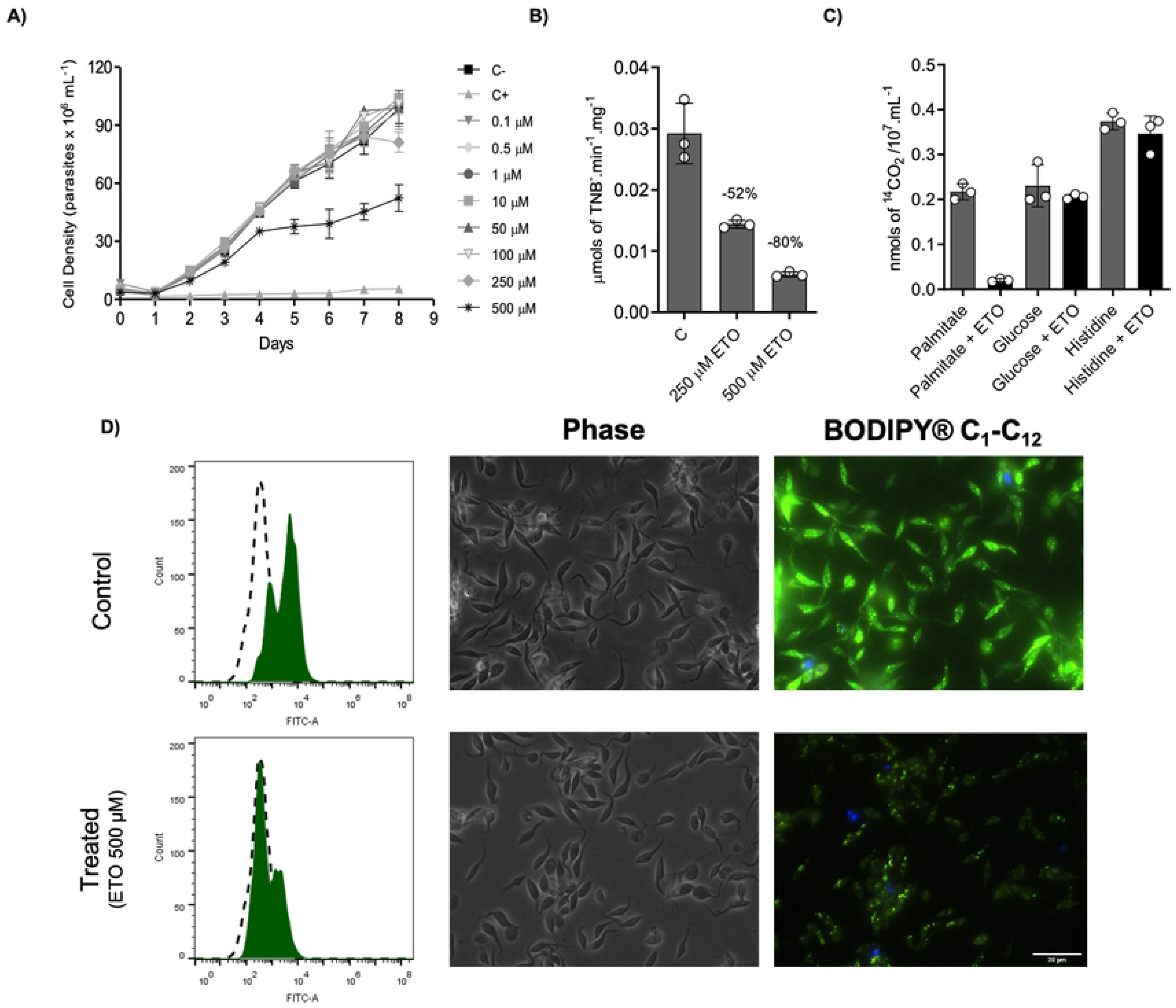
ETO inhibits CPT1 and interferes with cell proliferation in epimastigote forms. (A) Proliferation of epimastigote forms in the presence of 0.1 to 500 μM ETO. For the positive control of dead cells, a combination of antimycin (0.5 μM) and rotenone (60 μM) was used. (B) Inhibition of CPT1 activity in crude extracts using 250 and 500 μM of ETO. C) ^14^CO_2_ capture from ^14^C-U-palmitate oxidation. D) Flow cytometry analysis and fluorescence microscopy of epimastigote forms treated (or not) with ETO. In the histograms, dashed peaks represent unstained parasites and green-filled peaks represent parasites stained with BODIPY C_1_-C_12_. All the experiments were performed in triplicates. Statistical analysis was performed with one-way ANOVA followed by Tukey’s post-test at p < 0.05 using the GraphPad Prism 8.0.2 software program. We represent the level of statistical significance in this figure as follows: *** p value < 0.001; ** p value < 0.01; and * p value < 0.05. For p values > 0.05, we consider the differences not significant (ns).

### Etomoxir treatment affects cell cycle progression

The metabolic interference of ETO diminished epimastigote proliferation; however, this finding could be due to a decrease in the parasite proliferation rate or an increase in the death rate. Therefore, we checked if this compound could induce cell death through programmed cell death (PCD) or necrosis. PCD is characterized by biochemical and morphological events such as exposure to phosphatidylserine, DNA fragmentation, decreases (or increases) in the ATP levels, and increases in reactive oxygen species (ROS), among others [33]. The parasites were treated with 500 μM of ETO for 5 days, followed by incubation with propidium iodide (PI) for cell membrane integrity analysis and annexin-V FITC to evaluate the phosphatidylserine exposure. Parasites treated with ETO showed negative results for necrosis or programmed cell death markers (Fig 9A), indicating that the cell proliferation was arrested but cell viability was maintained. Because the multiplication rates seemed to be diminished, we performed a cell cycle analysis. Noticeably, the treated parasites were enriched in G1 (85.9%) with respect to non-treated cells (43.6%), suggesting that ETO prevented the entry of epimastigotes into the S phase of the cell cycle (Fig 9B). Last, we noticed that after washing out the ETO, the parasites recovered their proliferation at rates comparable to our untreated controls (Figs 9C).

**Figure 9.**
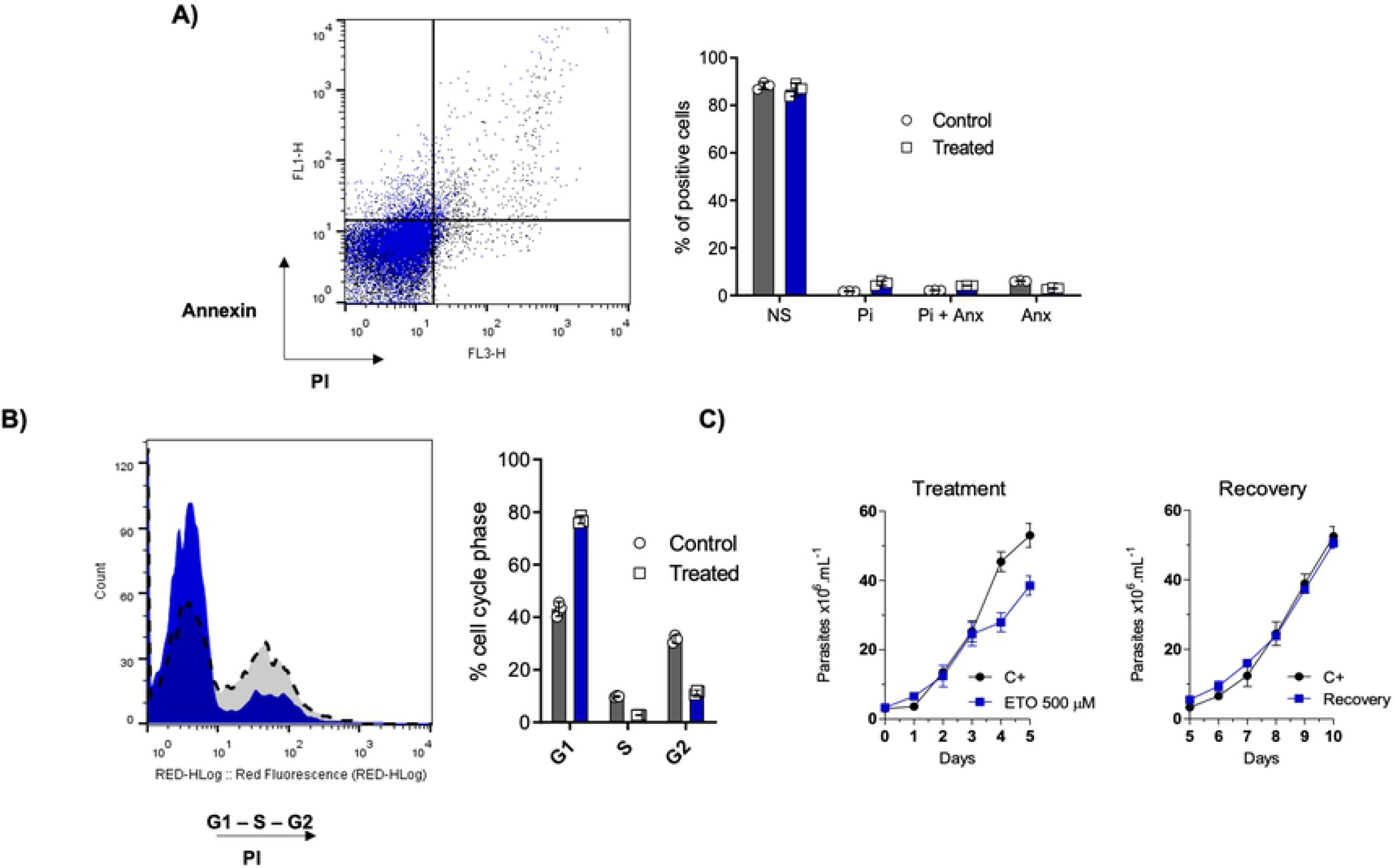
Analysis of extracellular phosphatidylserine exposure, membrane integrity and cell cycle after ETO treatment. Parasites in the exponential growth phase were treated with 500 μM of ETO for 5 days. (A) Following the incubation period, the parasites were labelled with propidium iodide (PI) and annexin V-FITC (ANX) and analysed by flow cytometry. (B) The cell cycle was assessed using PI staining. (C) Growth curves of epimastigote forms before and after removing the treatment. All the experiments were performed in triplicates. Statistical analysis was performed with one-way ANOVA followed by Tukey’s post-test p < 0.05, using the GraphPad Prism 8.0.2 software program. We represent the level of statistical significance in this figure as follows: *** p value < 0.001; ** p value < 0.01; and * p value < 0.05. For p values > 0.05, we consider the differences not significant (ns).

### Inhibition of FAO by ETO affects energy metabolism, impairing the consumption of endogenous fatty acids

The evidence obtained to date suggests that parasites resist metabolic stress by mobilizing and consuming stored fatty acids. Therefore, it is reasonable to hypothesize that ETO, which blocks the mobilization of fatty acids into the mitochondria for oxidation, probably perturbs the ATP levels in late-exponential or stationary phase cells. Parasites growing for 5 days under 500 μM ETO treatment or no treatment were collected to evaluate the ability of parasites that were treated or not with ETO to trigger oxygen consumption. The rates of O_2_ consumption corresponding to basal respiration were measured in cells resuspended in MCR respiration buffer. We then measured the leak respiration by inhibiting the ATP synthase with oligomycin A. Finally, to measure the maximum capacity of the electron transport system (ETS), we used the uncoupler FCCP [21]. Our results demonstrate that compared to no treatment, ETO treatment diminishes the rate of basal oxygen consumption, the leak respiration and the ETS capacity. In general, respiratory rates diminished in parasites treated with ETO when compared to the untreated ones. As expected, ETO treatment led to a 75% decrease in the levels of total intracellular ATP compared to untreated parasites (Fig 10A). To complement this result, because all these experiments were conducted in the complete absence of an oxidizable external metabolite, our results show that the parasite is able to oxidize internal metabolites (Figs 10B and 10C). Taking into account that treating parasites with ETO diminished the basal respiration rates of these parasites by approximately one-half (Figs 10B and 10C), it is reasonable to conclude that a relevant part of the respiration in the absence of external oxidisable metabolites is based on the consumption of internal lipids. This is consistent with the confirmation that epimastigotes maintain their viability in the presence of non-fatty acid carbon sources in the presence of ETO (S4 Fig). In summary, these results confirm that ETO is interfering with ATP synthesis through oxidative phosphorylation in epimastigote forms.

**Figure 10.**
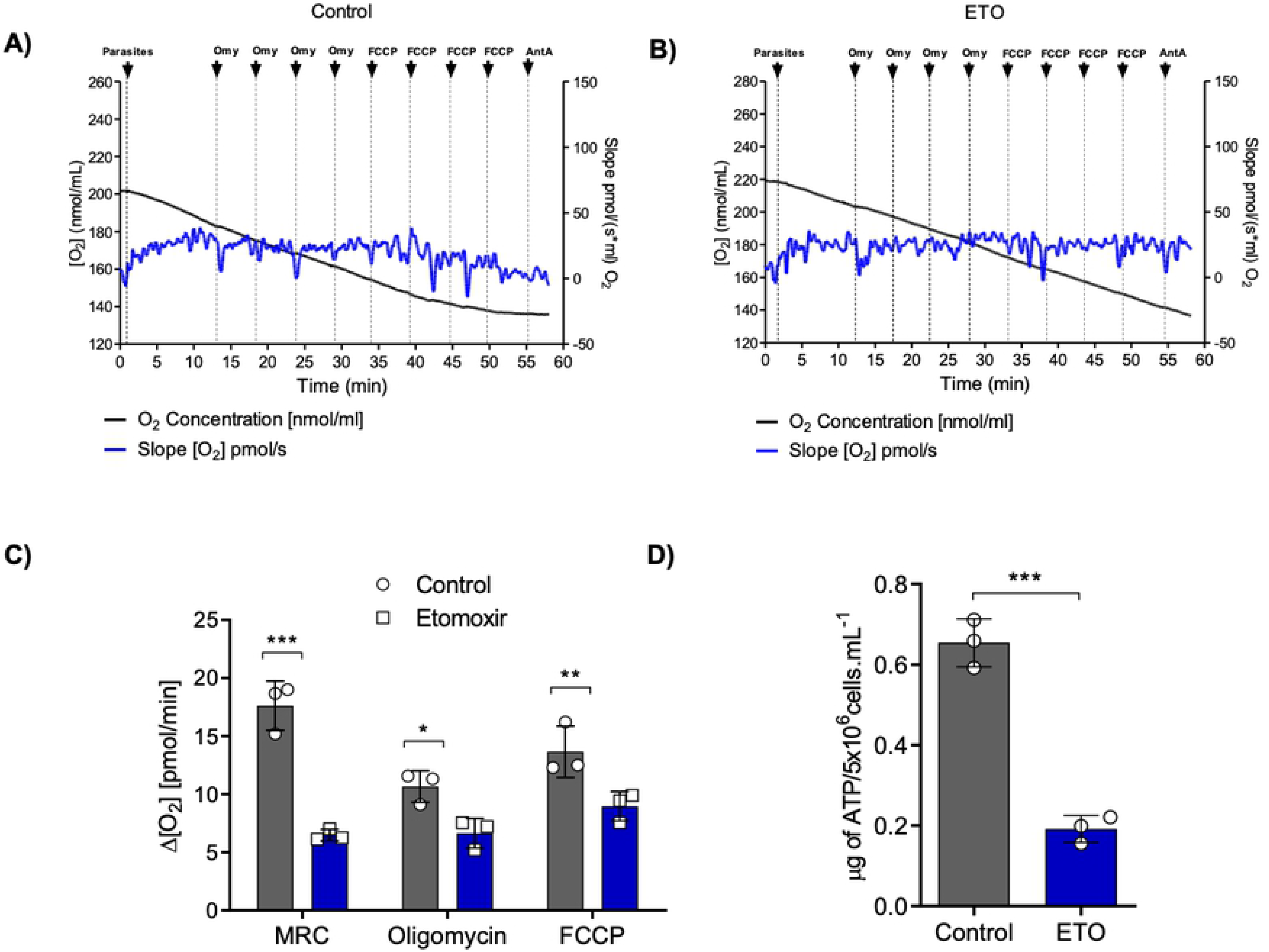
Effects of ETO on respiration and ATP production in epimastigote forms of T. cruzi. (A) Oxygen consumption of epimastigote forms after normal growth in LIT medium. (B) Oxygen consumption after ETO 500 μM treatment. Parasite growth in LIT medium with the compound until the 5^th^ day. In black, a time-course register of the concentration (pmols) of O_2_ in the respiration chamber. In blue, negative of the concentration derivative (pmols) of O_2_ with respect to time (velocity of O_2_ consumption in pmoles per second). The parasites were washed twice in PBS and kept in MRC buffer at 28 °C during the assays (see Materials and Methods for more details). (C) The basal respiration (initial oxygen flux values, MRC), respiration leak after the sequential addition of 0.5 μg/mL of oligomycin A (2 μg/mL), and electron transfer system (ETS) capacity after the sequential addition of 0.5 μM FCCP (2 μM) were measured for each condition. (D) Intracellular levels of ATP after treating with 500 μM ETO. The intracellular ATP content was assessed following incubation with different energy substrates or not (PBS, negative control). The ATP concentration was determined by luciferase assay and the data were adjusted by the number of cells. All the experiments were performed in triplicates. Statistical analysis was performed with one-way ANOVA followed by Tukey’s post-test at p < 0.05 using GraphPad Prism 8.0.2 software. We represent the level of statistical significance as follows: *** p value < 0.001; ** p value < 0.01; and * p value < 0.05. For p values > 0.05, we consider the differences not significant (ns).

### Endogenous fatty acids contribute to long-term starvation resistance in epimastigote forms

As previously demonstrated, ETO interferes with the consumption of endogenous fatty acids, and this impairment causes ATP depletion and cell cycle arrest. One intriguing characteristic of the insect stages of *T. cruzi* is their resistance to starvation. To observe the importance of internal fatty acids in this process, we incubated epimastigotes in PBS in the presence (or absence) of 500 μM ETO. The mitochondrial activity of these cells was followed for 24 h with Alamar blue®. Our results showed that the mitochondrial activity of the parasites in the presence of ETO was reduced by 31% after 48 h of starvation, and 65% after 72 h of starvation (Fig. 11) compared to the controls (untreated parasites). These data confirmed our hypothesis that the breakdown of accumulated fatty acids partially contributes to the resistance of the parasite under severe starvation.

**Figure 11.**
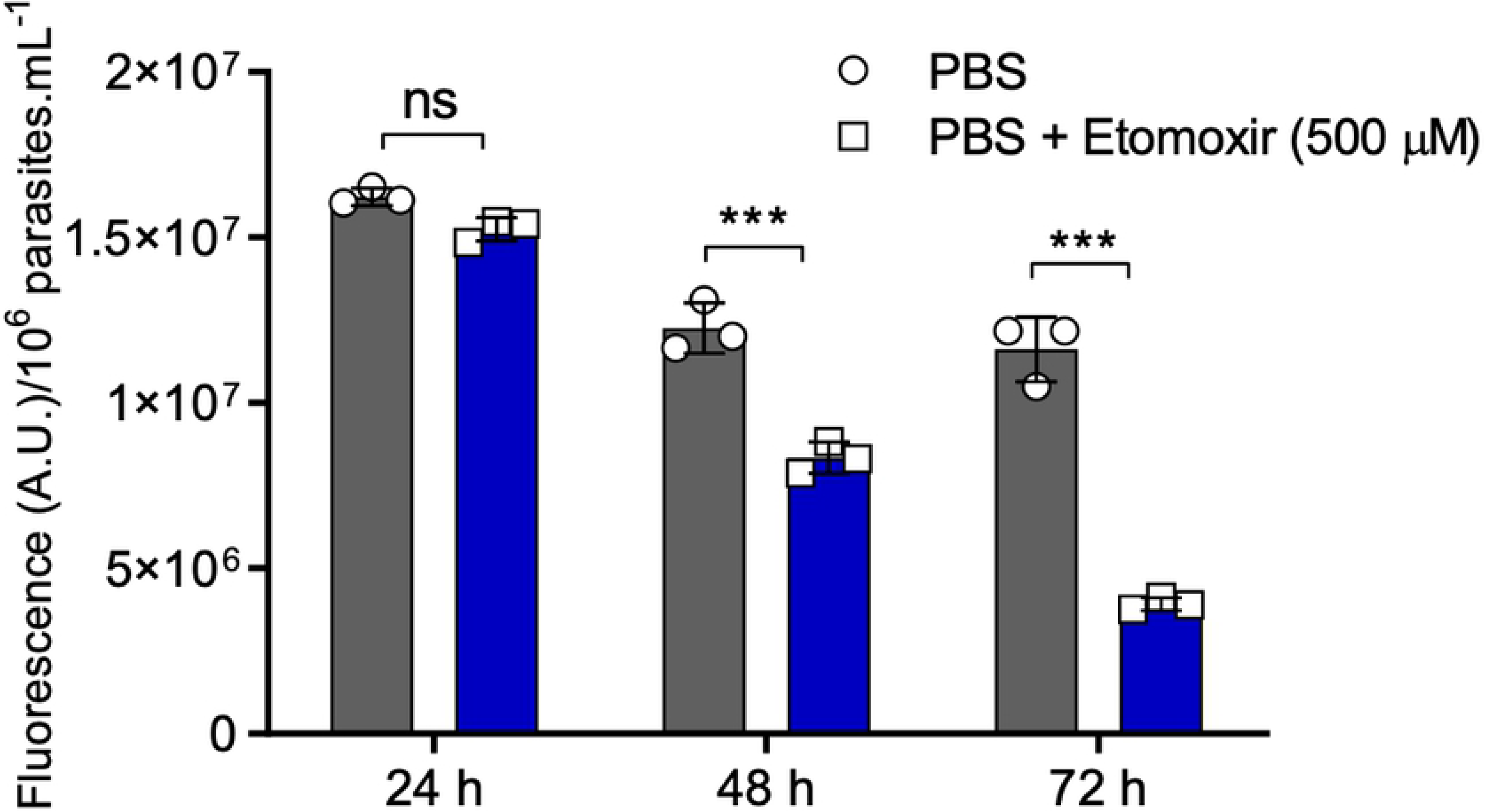
Internal fatty acid consumption contributes to parasite viability under severe nutritional starvation. Viability of epimastigote forms after incubation in PBS with or without ETO. The viability was assessed every 24 h using Alamar Blue®. Statistical analysis was performed with one-way ANOVA followed by Tukey’s post-test p < 0.05 using GraphPad Prism 8.0.2 software. We represent the levels of statistical significance as follow: *** p value < 0.001, and for p values > 0.05, we consider the differences not significant (ns).

### Inhibition of CPT1 impairs metacyclogenesis

Considering that the FAO increases in the epimastigotes during the stationary phase, and that differentiation into infective metacyclic trypomastigotes (metacyclogenesis) is triggered in the stationary phase of epimastigote parasites, one might expect a possible relationship between the consumption of fatty acids and metacyclogenesis. To approach this possibility, we initially compared the CPT1 activity of stationary epimastigote forms before and after a 24 h incubation in the differentiation medium TAU-3AAG. As observed, there is an increase in CPT1 activity after submitting the parasites to the metacyclogenesis *in vitro* (Fig. 12A). Parasites were then submitted to differentiation with TAU-3AAG medium in the presence of the probe BODIPY. The probe was incorporated into lipid droplets, confirming that fatty acids metabolism was active during the beginning of metacyclogenesis (Fig 12B). To address the importance of FAO during differentiation, metacyclogenesis was induced *in vitro* on ETO-treated or untreated (control) parasites. ETO treatment interfered with differentiation, diminishing the number of metacyclic forms present in the culture (Fig 12C). In addition, this inhibition was dose-dependent, with an IC_50_ = + 32.96 μM (Fig 12D). Importantly, we ruled out that the variation found in the differentiation rates was due to a selective death of treated epimastigotes, since their survival during this experiment in the presence or absence of ETO (from 5 to 500 μM) was not significantly different (**S5 Fig**). Based on these data, we could conclude that fatty acid oxidation, at the level of the CPT1, was also participating in the regulation of metacyclogenesis.

**Figure 12.**
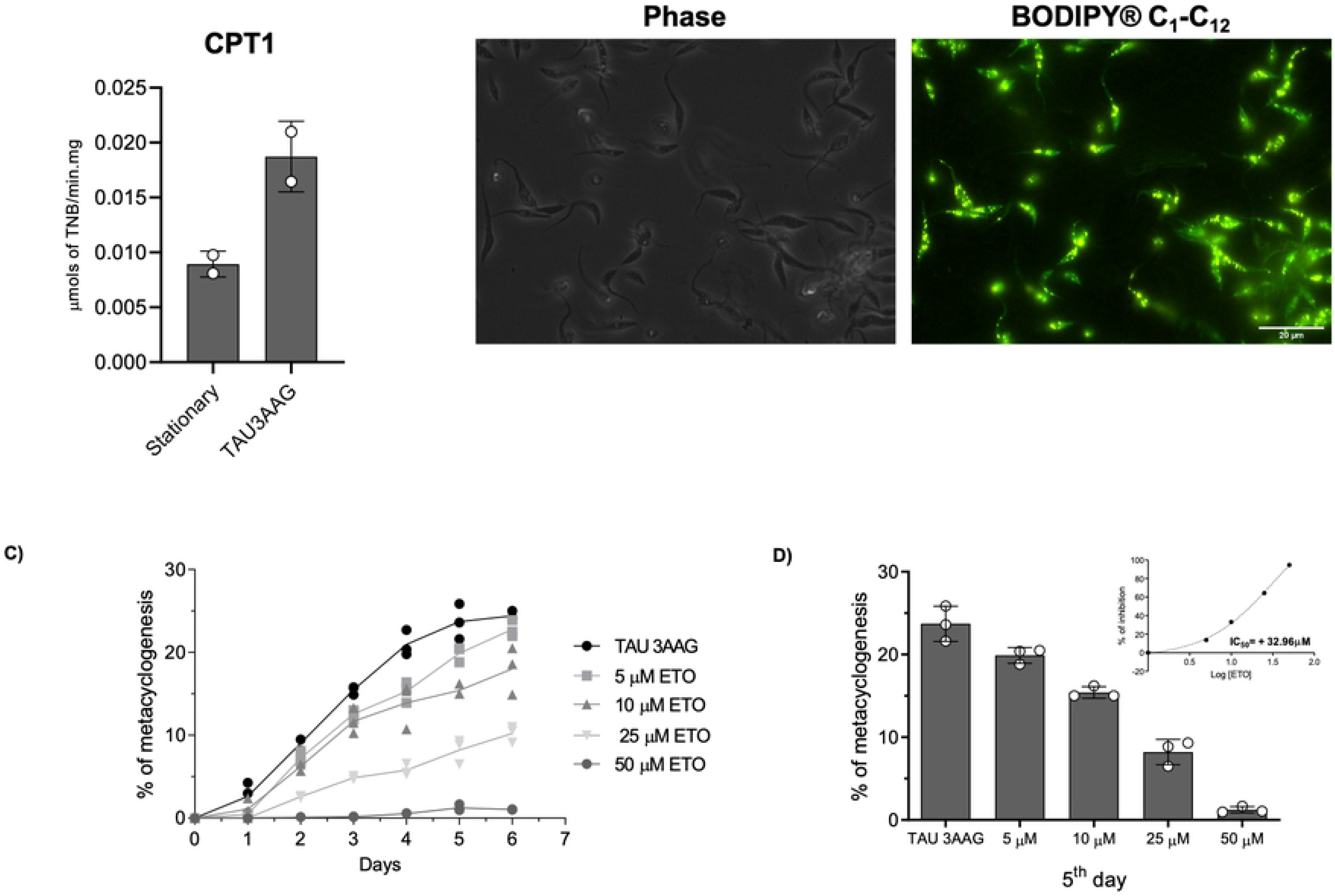
ETO inhibits metacyclogenesis. A) CPT1 activity of epimastigote forms in stationary phase and 24h after incubated in TAU-3AAG medium (for triggering metacyclogenesis). B) Fluorescence microscopy of cells incubated in TAU-3AAG in the presence of BODIPY® 500-510 C_1_-C_12_. C) Effects of different ETO concentrations on metacyclogenesis. The differentiation was evaluated by counting the cells in a Neubauer chamber each day for 6 days. This experiment was performed in triplicate. D) Percentage of differentiation at the 5^th^ day of differentiation. Inset: IC_50_ of metacyclogenesis inhibition by ETO. The enzymatic activities were measured in duplicate. All the other experiments were performed in triplicates.

## Discussion

During the journey of *T. cruzi* inside the insect vector, the glucose levels decrease rapidly after each blood meal [34], leaving the parasite exposed to an environment rich in amino and fatty acids in the digestive tube of *Rhodnius prolixus* [35,36]. Because the digestive tract of triatomine insects possesses a perimicrovillar membrane, which is composed primarily of lipids and is enriched by glycoproteins [37], it has been speculated that its degradation could provide lipids for parasite metabolism [38]. In this study, we showed that the insect stages of *T. cruzi* coordinate the activation of fatty acid consumption with the metabolism of glucose. Our experiments corroborate early studies about the relatively slow use of palmitate as an energy source by proliferating epimastigotes [39,40]. In addition, our results shed light on the end product excretion by epimastigote forms during incubation under starvation conditions, and during their recovery from starvation using glucose or palmitate. First, we showed that non-starved and starved parasites recovered in the presence of glucose, excreting succinate as their primary metabolic waste, as expected [41–43]. After 16 h of nutritional starvation, the consumption of internal carbon sources produces acetate as the primary end-product. In the presence of glucose after 16 h of starvation, we found that glucose-derived carbons contribute to the excreted pools of acetate and lactate. Interestingly, palmitate metabolism contributed to the increase in acetate production, followed by the production of alanine, pyruvate, succinate and lactate. The unexpected production of alanine, pyruvate and lactate can be explained by an increase in the TCA cycle activity, producing malate, which can be converted into pyruvate by the decarboxylative reaction of the malic enzyme (ME) [44]. Pyruvate can be converted into alanine through a transamination reaction by an alanine-[45], a tyrosine-[46] an aspartate aminotransferase [47], or a reductive amination by an alanine dehydrogenase [48]. The excretion of lactate could be a consequence of lactate dehydrogenase activity. However, it should be noted that this enzymatic activity has not been observed to date. In relation to the succinate production, a relevant factor favouring this process is the production of NADH by the third step of the beta-oxidation (3-hydroxyacyl-CoA dehydrogenase). This NADH can be oxidized through the activity of NADH-dependent mitochondrial fumarate reductase [49], which concomitantly converts NADH into NAD^+^ and fumarate into succinate. This succinate can be excreted or re-used by the TCA cycle, and the resulting NAD^+^ can be used as a cofactor for other enzymes.

As previously mentioned, it is well known that during the initial phase of proliferation, epimastigotes preferentially consume glucose, and during the stationary phase, a metabolic switch occurs towards the consumption of amino acids [8,10,42]. Our results show that this switch constitutes a broader and more systemic metabolic reprogramming, which also includes FAO. We detected this switch through changes in the enzymatic activities of key enzymes responsible for the regulation of FAO, such as CPT1 and ACC, which have increased and decreased activities, respectively, in the presence of glucose. Our findings showed that the inhibition of CPT1 affects the late phase of proliferation of epimastigotes when the switch to FAO has already occurred.

An interesting question about *T. cruzi* epimastigotes is how they survive long periods of starvation. Early data showed high respiration levels in epimastigotes incubated in the absence of external oxidisable carbon sources. This oxygen consumption was attributed to the breakdown of TAGs into free fatty acids and their further oxidation [50]. Here, we confirmed this finding by inhibiting the internal fatty acid consumption, which in turn diminished the oxidative phosphorylation activity, internal ATP levels and the total reductive activity of parasites under severe nutritional stress. Even more notably, we showed that under these conditions, the lipids stored in lipid droplets [51,52] are consumed. Unlike what has been observed in procyclic forms of *T. brucei*, in which the function of lipid droplets is not clear [53], our results show that in *T. cruzi*, they are committed to epimastigote survival under extreme metabolic stress. Of course, the contribution of other metabolic sources and processes such as autophagy in coping with nutritional stress cannot be ruled out [54].

Multiple metabolic factors has been involved in metacyclogenesis, such as the proline, aspartate, glutamate [55], glutamine [17] and lipids present in the triatomine digestive tract [56]. Interestingly, the occurrence of metacyclic trypomastigotes in culture leads to an increase in CO_2_ production from labelled palmitate [39]. The ETO treatment inhibited metacyclogenesis *in vitro*, showing that the consumption of internal fatty acids is important for cell differentiation. Consequently, we propose that lipids are not only external signals of metacyclogenesis, as previously suggested [56], but they also have a central role in the bioenergetics of metacyclogenesis. As in the oxidation of several amino acids, the acetyl-CoA produced from beta-oxidation and probably the reduced cofactors resulting from these processes are contributing to the mitochondrial ATP production necessary to support this differentiation step.

In conclusion, fatty acids are important carbon sources for *T. cruzi* epimastigotes in the absence of glucose. Palmitate can be taken up by the cells and fuel the TCA cycle by producing acetyl-CoA, the oxidation of which generates CO_2_. However, in the absence of external carbon sources, lipid droplets become the primary sources of fatty acids, helping the organism to survive nutritional stress. Importantly, FAO supports endogenous respiration rates and ATP production and powers metacyclogenesis.

## Acknowledgements

We thank the Core Facility for Scientific Research at the University of Sao Paulo (CEFAP-USP/FLUIR) for the flow cytometry analysis and Dr. Mauro Javier Veliz Cortez (Department of Parasitology, ICB-USP) for the microscopy work. We thank the Core Facility for Scientific Research at the University of Sao Paulo (CEFAP-USP/FLUIR) for the flow cytometry analysis and Dr. Mauro Javier Veliz Cortez (Department of Parasitology, ICB-USP) for the microscopy support.

## Supporting information

**S1**

**S1 Fig. ^1^H-NMR analysis of excreted end products from glucose and threonine metabolism.** The metabolic end products (succinate, acetate, alanine and lactate) excreted by the epimastigote cells that were incubated after 6 h in PBS (A), PBS after 16 h of starvation without (B) or with D-[U-^13^C]-glucose (C) or palmitate (D) were determined by ^1^H-NMR. Each spectrum corresponds to one representative experiment from a set of at least 3. A part of each spectrum ranging from 0.5 ppm to 4 ppm is shown. The resonances were assigned as indicated: A_12_, acetate; A_13_, ^13^C-enriched acetate; Al_12_, alanine; Al_13_, ^13^C-enriched alanine; G_13_, ^13^C-enriched glucose; L_12_, lactate; L_13_, ^13^C-enriched lactate; P_12_, palmitate; S_12_, succinate; and S_13_, ^13^C-enriched succinate.

**S2**

**S2 Fig. Time course activities of enzymes measured in this work.** A) (ACC) acetyl-CoA carboxylase, B) (CPT1) carnitine-palmitoyltransferase, and C) (SPT) serine palmitoyltransferase. All the activities were measured in cell-free extracts of epimastigote forms at different moments of the growth curve as indicated in the main text. All the measurements were performed in triplicates.

**S3**

To check if other well-known FAO inhibitors have the same effect on the proliferation of *T. cruzi* epimastigotes, we performed the same assay as described in Materials and Methods by evaluating different concentrations of valproic acid (AV) [57], trimetazidine [58,59] and β-hydroxybutyrate [60], which are inhibitors of 3-ketothiolase. Because they did not affect the proliferation of the epimastigote forms, we used the higher concentration evaluated in these assays to know if the compounds inhibit FAO by ^14^CO_2_ trapping by using U-^14^C-palmitate as a substrate. As observed, none of these compound inhibited the ^14^CO_2_ production from palmitate, confirming that they are not inhibiting FAO in *T. cruzi*.

**S3 Fig. Other FAO inhibitors did not affect cell proliferation and FAO in the epimastigote forms.** The compounds were evaluated at concentrations between 0.1 and 1000 μM. For positive controls of dead cells, a combination of antimycin (0.5 μM) and rotenone (60 μM) were used. The maximum concentration tested for these compounds does not diminish CO_2_ liberation from FAO. A) Valproic Acid (AV). B) Trimetazidine (TMZ). C) β-hydroxybutyrate (βHOB).

**S4**

In this study, we showed that the epimastigote forms of *T. cruzi* present low sensitivity in response to ETO treatment. Recently, some groups described off-target effects when ETO is used at concentrations of up to 200 μM [61,62]. To validate ETO as an FAO inhibitor of *T. cruzi*, the parasites were incubated for 24 h in PBS (negative control), 0.1 mM palmitate supplemented with BSA, 5.0 mM histidine, 5 mM glucose, 0.1 mM carnitine and BSA without adding palmitate in the presence (or not) of 500 μM ETO. The viability of these cells was inferred from the measured total reductive activity using MTT assays (see Material and Methods section for more details). As expected, ETO treatment did not affect the viability of cells incubated in glucose or histidine but did affect the viability of the cells incubated with palmitate or carnitine. Surprisingly, we also observed an ETO effect on parasites under metabolic stress, such as those incubated with PBS or BSA. This finding could be explained by the fact that under metabolic stress, the parasite mobilizes and consumes its internal lipids.

**S4 Fig. ETO did not affect the viability of epimastigote forms in the presence of other carbon sources.** The viability of epimastigote forms after incubation with different carbon sources and palmitate. The viability was assessed after 24 h using MTT.

**S5**

Because metacyclogenesis occurs in chemically defined conditions, we performed a viability assay to define the maximum tolerated concentration that allows the parasites to survive under ETO treatment. Stationary epimastigotes in TAU-3AAG media were treated with different concentrations of ETO (range 5 to 500 μM) during 24 h. The viability of these cells was inferred by measuring the total reductive activity using an Alamar blue assay [63]. Briefly, after 24 h in the presence or absence of ETO, the cells were incubated with 0,125 μg.mL^−1^ of Alamar blue reagent in accordance with the protocol by [17]. Under these conditions, the parasites were 10 times more sensitive to ETO treatment, surviving when subjected to ETO concentrations between 5-50 μM (Fig. S3 A). This range of concentrations used to treat the parasites was maintained in TAU-3AAG medium and to follow the differentiation by daily counts, based on the percentage of metacyclic trypomastigotes collected in culture supernatant. To confirm that the parasites were still alive after 5 days under differentiation, we checked the viability of cells that were treated (or not, control) using the same assay. As shown above (Figure S3 B), the parasites were viable under all the tested conditions. Considering that TAU-3AAG contains glucose in its composition, we performed an *in vitro* metacyclogenesis using only proline as a metabolic inducer [64]. As observed, even in the absence of glucose, ETO treatment affects metacyclogenesis.

**Fig S5. Viability of epimastigote forms subjected to metacyclogenesis under different ETO concentrations.** A) Cell viability under metacyclogenesis after 24 h of treatment with different ETO concentrations. B) Cell viability under metacyclogenesis after 5 days in the presence of ETO. C) Effect of ETO on the metacyclogenesis induced by proline.

